# Serotonergic control in initiating defensive responses to unexpected tactile stimuli in the trap-jaw ant *Odontomachus kuroiwae*

**DOI:** 10.1101/2020.04.28.065466

**Authors:** Hitoshi Aonuma

**Author notes:** Correspondence to be sent to: Hitoshi Aonuma, Research Institute for Electronic, Science, Hokkaido University, Sapporo, Hokkaido, 060-0812, Japan.

## Abstract

A decision to express a defensive response or an escape response to a potential threat is crucial for insects to survive. This study investigated an aminergic mechanism underlying defensive responses to unexpected touch in an ant that has powerful mandibles, the so-called trap-jaw. The mandibles close extremely quickly and are used as a weapon during hunting. Tactile stimulation to the abdomen elicited quick forward movements in a “dart escape” in 90% of the ants in a colony. Less than 10% of the ants responded with a quick “defensive turn” towards the source of stimulation. To reveal the neuronal control mechanisms underlying this defensive behavior, the effects of brain biogenic amines on the responses to tactile stimulus were investigated. The levels of octopamine (OA), dopamine (DA) and serotonin (5HT) in the brain were significantly elevated in ants that responded with a defensive turn to the unexpected stimulus compared to ants that responded with a dart escape. Oral administration of DA and 5HT demonstrated that both amines contributed to the initiation of a defensive response to the stimulus. Oral administration of L-DOPA weakly affected the initiation of the defensive turn, while 5HTP strongly affected the initiation of defensive behavior. Oral administration of an antagonist of 5HT, ketanserin, abolished the effect of 5HTP. These results indicate that endogenous 5HT in the brain has a key role to play in modulating the initiation of defensive behavior in the trap-jaw ant.

## INTRODUCTION

Ponerine ants hunt and consume small arthropods that represent the major component of their diet (Choe and Crespi, 1997). During hunting forager ants must increase their levels of aggressiveness to attack and capture prey, and to defend against unexpected encounters with an enemy. Defensive behavior can escalate to violent attacks against opponents, which also increases the risk of damage. To avoid such risks, the choice to escape could increase survivability. Similar to individual behavior, social decisions to defend or escape are crucial for a colony to survive (Holway et al., 1998).

The ant genus *Odontomachus* have long and powerful mandibles, the so-called trap-jaw, that functions as a weapon when hunting (De la Mora et al., 2008). The trap-jaw ant captures prey by closing the mandibles extremely quickly (S.1) (Gronenberg, 1996a; Just and Gronenberg, 1999). If the ant encounters a potential threat, however, the ant responds rapidly with a defensive turn response or with a dart escape response (S. 2). Behavioral responses with both defensive turns and dart escape to unexpected tactile stimulation are common to other arthropod such as crayfish (Nagayama et al., 1986) and crickets (Alexander, 1961). It is believed that social experience influences the behavioral response to an unexpected tactile stimulus (Song et al., 2006).

Biogenic amines function as neurotransmitters, neuromodulators and neurohormones in the brain and play key roles in behavior (Evans, 1980; Baumann et al., 2003; Roeder, 2005). The octopaminergic (OAergic) system is thought to be associated with nest-mate recognition in social insects such as honeybees (Robinson et al., 1999) and ants (Vander Meer et al., 2008). Aminergic systems in the brain are closely associated with aggressive behavior in arthropods (Edwards and Kravitzt, 1997; Kravitz, 2000; Stevenson et al., 2005; Hoyer et al., 2008); Johnson et al., 2009; Rillich et al., 2011a). The effects of biogenic amines on aggressive behavior have also been demonstrated in ants (OA: (Aonuma and Watanabe, 2012b; Yakovlev, 2018), TA: (Szczuka et al., 2013), DA: (Ohkawara and Aonuma, 2016; Vander Meer et al., 2008) (Shimoji et al., 2017), 5HT: (Kostowski et al., 1975). This study focuses on the behavioral responses to unexpected tactile stimuli to gain an understanding of the neuronal mechanisms underlying the initiation of defensive behavior in the trap-jaw ant. Moreover, the study asks how biogenic amines contribute to initiate behavioral responses to such unexpected tactile stimuli, since social interactions between nest-mates influences aminergic homeostasis in the brain (Wada-Katsumata et al., 2011).

The levels of brain amines of the workers were measured and compared between those that respond with a defensive turn and those that respond with an escape dart to tactile stimulation. The effects of oral administration of amines on initiating defensive behavior were then examined to confirm whether brain amines are associated with initiating defensive behavior. This work provides insights into understanding how trap-jaw ants initiate defensive behavior against an unexpected encounter with an enemy.

## MATERIALS AND METHODS

### Animals

Workers of the trap jaw ants *Odontomachus kuroiwae* were used throughout this study. Colonies of ants were collected in Okinawa, Japan. They are mostly polygynous and contained 3-4 queens, 200-300 workers and broods. Each colony was installed into an artificial plaster nest (2000mm ×2000mm×40mm) in a plastic case (6000mm ×4000mm×2000mm) on a 14h:10h light and dark cycle (lights on at 6:00) at 25±2°C. Ants were fed a diet of insect jelly (Marukan Co., Ltd, Osaka, Japan), cricket nymphs and water *ad libitum*. There are no distinctive differences between major and minor workers in *O. kuroiwae*.

### Behavior experiments

*O. kuroiwae* workers were randomly collected from colonies and kept isolated in a plastic petri dish (φ50mm) for 60min before behavioral experiments. To elicit defensive turns or dart escape behavior, the abdomen of an ant was touched gently using the tip of a fine paint brush. A similar behavioral assay was used in previous studies on crayfish (Nagayama et al., 1986; Aonuma et al., 1994). Behavioral responses to the tactile stimulation of the abdomen were observed and recorded using a digital video camera (JVC, GC-P100, Tokyo Japan) for later analysis. The defensive levels were classified into 4 types (Fig. 1, S. 2). An ant that responded with a quick forward movement, the so-called *dart escape*, to a tactile stimulus was scored as “*Level -1*”. If an ant did not respond to the stimulus, it was scored as “*Level 0*”. Defensive turn responses were divided into 2 types so that if an ant turned towards the stimulus without opening its mandibles it was scored as “*Level 1*”, while if it turned with its mandibles open it was scored as “Level 2”.

**Fig. 1.**
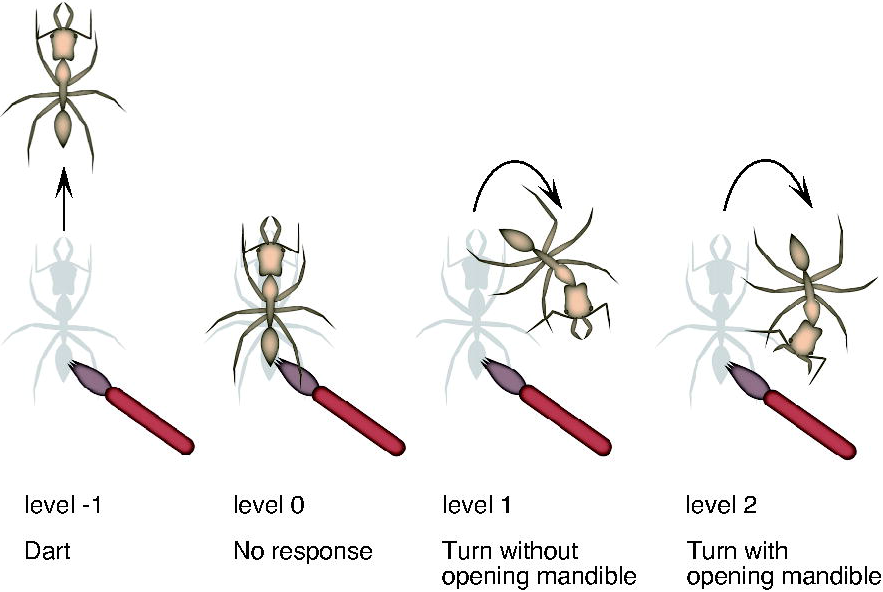
Pictograms illustrating the responses to tactile stimulation of the abdomen of the ants. The response was classified into 4 levels (level -1 to 2). *Level -1* dart response: the ant quickly moved forward. *Level 0* no response: the ant shows no obvious response to a tactile stimulus. *Level 1* turn responses without opening of the mandibles: the ant orientated towards the source of the stimulus without opening the mandibles. *Level 2* turn responses with open mandibles: the ant orientated towards the source of the stimulus and opened the mandibles widely.

### Measurement of brain biogenic amines

The levels of biogenic amines in the brain were measured using high-performance liquid chromatography (HPLC) with electrochemical detection (ECD) (Wada-Katsumata et al., 2011; Aonuma and Watanabe, 2012a). Workers of *O. kuroiwae* whose body mass were 9.9 ± 1.1mg (N = 60, mean ± SD) were randomly collected from colonies. Since the levels and content of brain amines change depending on age (Seid and Traniello, 2005; Seid and Traniello, 2006; Aonuma and Watanabe, 2012a), newly emerged workers were not used in this study. Test animals were sampled between 10:00 -12:00 to avoid circadian effects (Tomioka et al., 1993). This study compared the levels of brain amines of ants that showed dart responses (*level -1*) and turn responses (*level 2*) to tactile stimulation of the abdomen. Thirty *level -1* workers and another 30 *level 2* workers were collected from 3 different colonies (10 ants from each colony). Ants were then quickly frozen in liquid N_2_ to inhibit enzyme activity. A brain of an ant was dissected out in ice-cold normal saline (128.3 mM NaCl, 4.7 mM KCl, 1.63 mM CaCl_2_, 6 mM NaHCO_3_, 0.32 mM NaH_2_PO_4_, 82.8 mM trehalose, pH 7.4). Each brain was placed into a micro glass homogenizer and homogenized with 50μl of ice-cold 0.1M perchloric acid containing 5ng of 3, 4-dihydroxybenzylamine (DHBA, SIGMA, St Louis, MO, USA) as an internal standard. After centrifugation of the homogenate (0°C, 15000g, 30min), 35μl of supernatant was collected.

The HPLC-ECD system comprised a pump (EP-300, EICOM Co., Kyoto, Japan), an auto-sample injector (M-504, EICOM Co., Kyoto, Japan) and a C18 reversed-phase column (250 mm × 4.6 mm internal diameter, 5μm average particle size, CAPCELL PAK C18MG, Shiseido, Tokyo, Japan) heated to 30 °C in the column oven. A glass carbon electrode (WE-GC, EICOM Co.) was used for electrochemical detection (ECD-100, EICOM Co.). The detector potential was set at 890mV versus an Ag/AgCl reference electrode, which was also maintained at 30°C in a column oven. The mobile phase containing 0.18M chloroacetic acid and 16μM disodium EDTA was adjusted to pH 3.6 with NaOH. Sodium-1-octanesulfonate at 1.85mM as an ion-pair reagent and CH_3_CN at 8.40% (v/v) as an organic modifier were added into the mobile phase solution. The flow rate was kept at 0.7ml/min. The chromatographs were acquired using a computer program PowerChrom (eDAQ Pty Ltd, Denistone East, NSW, Australia). The supernatants of samples were injected directly onto the HPLC column. After acquisition, they were processed to obtain the amount of biogenic amines in the same sample by the ratio of the peak area of substances to the internal standard DHBA. A standard mixture for quantitative determination that contained amines, precursors and metabolites was used. Twenty compounds at 100ng/ml each were DL-3,4-Dihydroxy mandelic acid (DOMA), L-β-3,4-Dihydroxyphenylalanine (DOPA), L-Tyrosin (Tyr), N-Acetyloctopamine (Nac-OA), (-)-noradrenaline (NA), 5-Hydroxy-L-tryptophan (5-HTP), (-)-adrenaline (A), DL-Octopamine (OA), 3,4-Dihydroxybenzylamine (DHBA, as an internal standard), 3,4-Dihydroxy phenylacetic acid (DOPAC), N-Acetyldopamine (Nac-DA), 3,4-Dihydroxyphenethylamine (DA), 5-Hydroxyindole-3-acetic acid (5HIAA), N-Acetyltyramine (Nac-TA), N-Acetyl-5-hydroxytryptamine (Nac-5HT), Tyramine (TA), L-Tryptophan (Trp), 3-Methoxytyramine (3MTA), 5-Hydroxytryptamine (5HT), 6-Hydroxymelatonin (6HM). Nac-OA Nac-DA and Nac-TA were synthesized by Dr. Matsuo (Keio University, Japan). All other substances were purchased from Sigma-Aldrich (St. Louis, MO, USA).

### Pharmacological experiments

Ants were randomly collected from colonies and placed in plastic petri dishes. After each ant was isolated for 60min to rest, the response to a tactile stimulus was observed. Ants that showed a dart response to the tactile stimulus were used for pharmacological experiments. Pharmacological agents were dissolved in 20% sucrose solution and orally applied to the ants. For control a 20% sucrose solution was used. To manipulate the levels of biogenic amines in the brain, serotonin (5HT), octopamine (OA), dopamine (DA), precursor of serotonin 5-Hydroxy-L-tryptophan (5HTP) and precursor of dopamine L-β-3,4-Dihydroxyphenylalanine (L-DOPA) were used. After oral administration of each agent, responses of the ant to the tactile stimulus were observed. To inhibit 5HT receptors, ketanserin was used (Vleugels et al., 2015). All substances were purchased from Sigma-Aldrich (St. Louis, MO, USA).

### Statistical analyses

Statistical analyses were performed using Graphpad Prism (Graphpad, v.8.4.2). A Kruskal-Wallis test was used to analyse the difference in behavioral responses among colonies. Differences in the levels of biogenic mines were tested using unpaired t tests with Whelch’s correction. Differences were considered significant at p<0.05 level (two-tailed). Analysis of variance (ANOVA) with Tukey’s multiple comparison test was used to analyse pharmacological experiments.

## RESULTS

### Dart escape and defensive turns

Trap-jaw ant *O. kuroiwae* workers were collected from 7 different colonies and behavioral responses to the tactile stimulus examined. In total 580 workers were randomly collected from different colonies and responses to the tactile stimulus observed. There was no significant difference among colonies in responses of the ants to the stimulus (Kruskal-Wallis test). Most of workers (523 in total out of 580 ants from 7 colonies, 90.2±5.2%, Mean ± SD) showed dart escape response (*level -1*) to the stimulus. As soon as the tip of a fine drawing brush made contact with the abdomen, the ants quickly moved forward away from the stimulus source. 2.4 ±1.4% (14 in total out of 580) of the ants showed no obvious response to the stimulus *(level 0*). The behavior of *level 0* ants was clearly different from *level -1* dart responses as the ants did not move away from the stimulus source. Even if the ants walked after the tactile stimulus, walking speed was much slower than that of dart escape response (S. 2). Approximately 10% of the ants responded with defensive turn to the tactile stimulus (including *level 1 and level 2*). The ants that responded with a *level 1* turn was 2.4±2.2% (14 in total out of 580 ants) and most of the ants showed antennal boxing towards the stimulus source after they turned towards it. 7.4±4.8% (29 in total out of 580 ants) of the ants responded with a *level 2* turn to the stimulus. These ants also showed antennal boxing toward the stimulus. Antennal boxing functions to detect and identify the source of unexpected stimulus and it has been reported in ants that antennal boxing is observed in dominant workers during agonistic interactions (Gobin et al., 2001). This indicates that aggressive workers showed turn responses to the tactile stimulus. However, there were few ants that escalated their aggressiveness to initiate violent attacks against the stimulus.

### Levels of biogenic amines in the brain

Thirty *level 2* workers that showed a turn response and 30 *level -1* workers that showed a dart response were collected from 3 different colonies (10 ants each from each colony), and biogenic amines in the brain analyzed using HPLC-ECD.

OA is generated from L-tyrosine through a different synthetic pathway from DA generation. Tyrosine decarboxylase generates TA and then tyramine β-hydroxylase generates OA. Both TA and OA were detected in the brains of all ants that showed both turn and dart responses. There was no significant difference in the amount of TA in the brains between the ants that showed *level 2* turn responses (0.42 ± 0.37 pmol/brain, N = 30, Mean ± SD) and *level -1* dart responses (0.36 ± 0.19 pmol/brain, N = 30) (Fig. 2A). On the other hand, the amount of OA in the brains of ants that showed *level 2* turn responses (0.94 ± 0.28 pmol/brain, N = 30) was significantly greater (p = 0.0226) than in the ants that showed *level -1* dart response (0.76 ± 0.29 pmol/brain, N = 30) (Fig. 2B). Nac-OA and NacTA are catabolites of OA and TA, respectively, by the activation of arylalkylamine N-acetyltransferase. This study failed to detect Nac-OA. Nac-TA was detected in the brains of all ants. There was also no significant difference in the levels of brain Nac-TA between ants that showed dart and turn responses (S.3A).

**Fig. 2.**
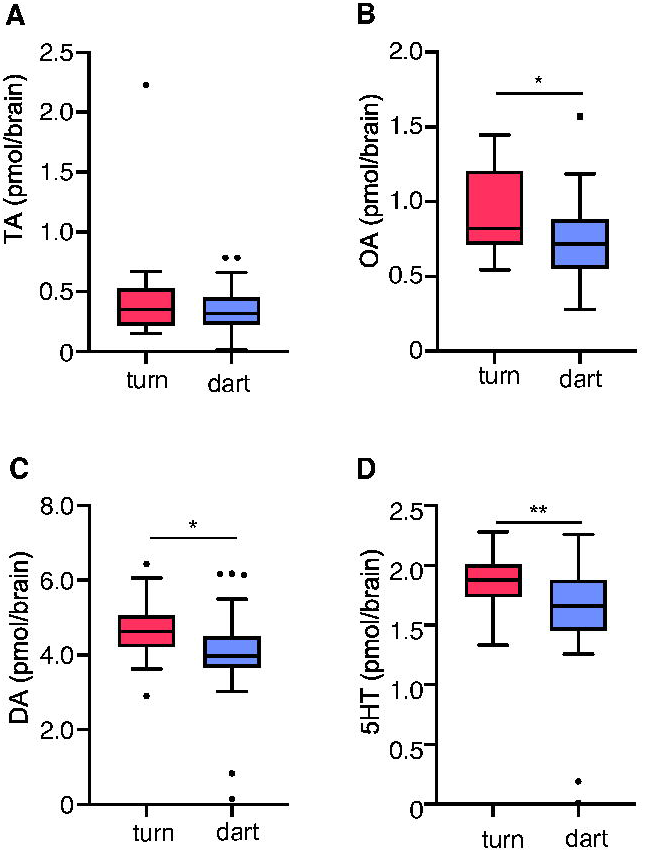
Contents of biogenic amines in the brains of ants that showed *Level 2* turn responses and those that showed *level -1* dart responses, to tactile stimulation. Box-and-whisker graphs indicates minimum, median, maximum, 25% percentile and 75% percentile. **A:** Content of TA in the brain. There was no significant difference in the contents of TA between the ants that showed *level 2* turn responses and those that showed *level -1* dart responses (p = 0.394). **B:** Content of OA in the brain. Brain OA levels in the ants that showed *level 2* turn response were significantly higher than in the ants that showed *level -1* dart responses (p = 0.0226). **C:** Content of DA in the brain. Brain DA levels in the ants that showed *level 2* turn responses were significantly higher than in the ants that showed *level -1* dart responses (p = 0.0329). **D:** Content of 5HT in the brain. Brain 5HT levels in the ants that showed *level 2* turn responses were significantly higher than in the ants that showed *level -1* dart responses (p = 0.0086).

DA is generated from DOPA by activation of aromatic L-amino acid decarboxylase and catabolized to Nac-DA by activation of arylalkylamine N-acetyltransferase. DA and Nac-DA were detected in all samples. The content of DA in the ants that showed *level 2* turn responses was 4.67 ± 0.73 pmol/brain (N = 30). The content of DA in the brains of ants that showed *level -1* dart responses was 4.07 ± 1.30 pmol/brain (N = 30). The content of DA in the brains of ants that showed turn responses was significantly elevated compared to ants that showed dart response (p = 0.0329) (Fig. 2C). DOPA is generated from tyrosine by the activation of tyrosine hydroxylase. This study failed to measure both tyrosine and DOPA since the peaks of these two substances on the chromatogram appeared with unknown front peaks. The catabolite of DA was detected in the brain. The amount of Nac-DA in the brains of *level 2* ants was 0.96 ± 0.43 pmol/brain (N=30) while that of *level -1* ants was 0.93 ± 0.36 pmol/brain (N = 30). There was no significant difference between these ants (S. 3B).

5HT was also detected in the brains of each ant. The level of 5HT in the brains of *level 2* ants was 1.87 ± 0.23 pmol/brain (N = 30) while that in *level -1* ants was 1.60 ± 0.48 pmol/brain (N = 30) (Fig. 2D). The levels of 5HT in the brain were elevated significantly in *level 2* ants compared to *level -1* ants (p = 0.0086). 5HT is generated from 5HTP by the activation of aromatic L-amino acid decarboxylase. 5HTP was also detected in the brains of each ant. The content of 5HTP in the brains of *level 2* ants was 0.34 ± 0.33 pmol/brain (N = 30) and slightly more than that in *level -1* ants (0.23 ± 0.34 pmol/brain, N = 30), although there was no statistically significant difference (S. 3C). Endogenous 5HT is catabolized to Nac-5HT by arylalkylamine N-acetyltransferase. Nac-5HT was detected at a level of 0.36 ± 0.22 pmol/brain (N = 30) in the brains of *level 2* ants while that in *level -1* ants was 0.30 ± 0.13 pmol/brain (N= 30) (S. 3D).

### Effects of oral administration of biogenic amines

Measurements of brain amines demonstrated that the levels of OA, DA and 5HT in ants that showed a turn response (*level 2*) to tactile stimulation were significantly higher than in ants that showed a dart response (*level -1*). To investigate which brain amines were associated with initiating defensive behavior, oral administration of the agonists was performed.

For control experiments, the effect of oral administration of a 20% sucrose solution was examined first (Fig. 3A). Sixty workers were randomly collected from 3 colonies (20 workers from each colony) and responses to the tactile stimulus observed. Fifty-four out of 60 ants responded with a dart escape, and these 54 ants were used to examine the effect of 20% sucrose solution on the response to the tactile stimulus. There was no significant change in the defensiveness level score after administration of 20% sucrose solution with most still responding with dart escape after 1hr and after 2hr.

**Fig. 3.**
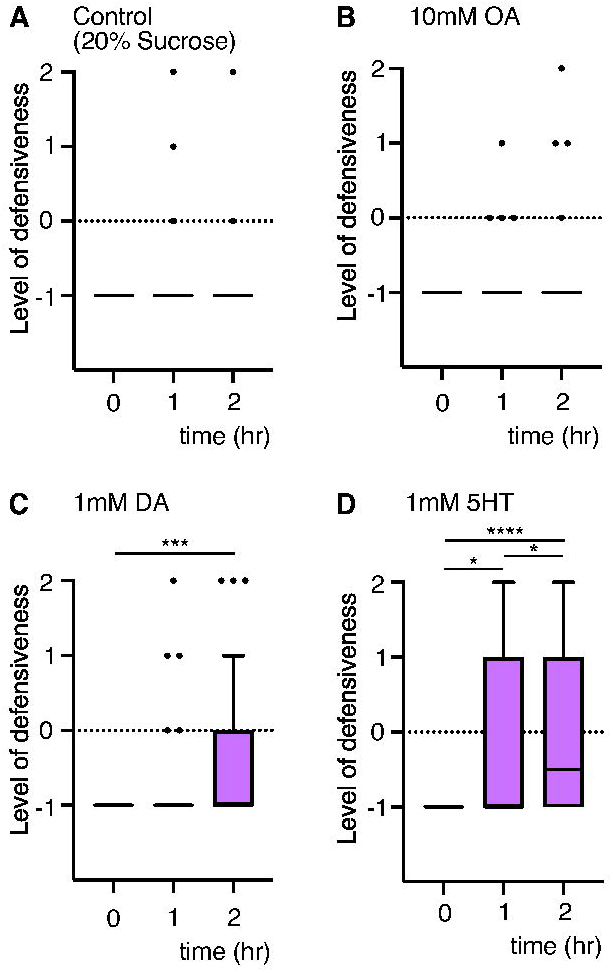
Pharmacological manipulation of biogenic amines in the brain. Box-and-whisker graphs indicate minimum, median, maximum, 25% percentile and 75% percentile. **A:** Effects of oral application of 20% sucrose solution on defensive behavior. The responses of the ants that showed dart responses to a tactile stimulus were examined (N = 54, collected from 3 different colonies). No significant changes in response to the tactile stimulus were observed after 1hr and after 2hr of application. **B:** Oral administration of 10mM OA (N = 35, collected from 2 different colonies). The ants that showed dart responses to the tactile stimulus did not initiate a turn response to the tactile stimulus after 1hr and 2hr. **C:** Oral administration of 1mM DA (N = 30 collected from 2 different colonies). Defensiveness level score were significantly increased after 2hr. **D:** Oral administration of 1mM 5HT (N=38, collected from 2 different colonies). Defensiveness level score increased significantly after 1hr and 2hr. *: p < 0.05, ***: p < 0.001, ****: p < 0.0001

Oral administration of OA did not significantly change the defensiveness level of the ants. Twenty workers were randomly collected from a colony and responses to tactile stimulation observed prior to the application of 1mM OA solution. All ants responded with a dart escape to the stimulus. Oral administration of 1mM OA did not change the response to the stimulus even after 2hr. The effect of a 10 mM OA solution on the behavior was then examined. Forty workers were randomly collected from 2 different colonies (20 ants from each) and the response to the tactile stimuli prior to oral administration of OA observed. Thirty-four out of 40 ants that received orally a 10mM OA solution responded with a dart escape in response to the tactile stimulus (Fig. 3B).

Most of these ants did not change their response to the stimulus. Three ants out of 34 showed no obvious response to the stimulus, and only 1 ant showed a turn response after 1hr. Four ants out of 34 showed turn response to the stimulus 2hr after administration (*level 0*: N = 1, *level 1*: 2, *level 2*: N = 1).

Oral administration of DA significantly increased the defensiveness level score of the ants (Fig. 3C). Thirty ants were randomly collected from 2 colonies (15 ants from each of 2 colonies) and responses to the tactile stimulus prior to the administration observed. All of them responded with a dart escape to the stimulus. Oral administration of 1mM DA significantly increased the defensiveness level score after 2hr (p = 0.0006), although there was no significant difference to that after 1hr (Fig. 3C). Three ants out of 30 responded with a turn behavior after 1hr following oral administration of DA (*level 0*: N=1, *level 1*: N = 2, *level 2*: N = 1). The number of ants that showed a turn response increased to 5 (*level 1*: N = 2, *level 2*: N = 3) and number of the ants that ignored the stimulus (*level 0*) was 9.

Oral administration of 1mM 5HT solution significantly increased the defensiveness level score after 1hr and 2hr (Fig. 3D). Forty workers were randomly collected from colonies (20 ants from each of 2 colonies) and responses to the tactile stimulus observed. Thirty-eight out of 40 ants showed a dart response to the stimulus and then used to examine the effect of administration of 5HT on the behavior. Ten out of 38 ants responded with a turn response to the tactile stimulus after 1hr of 1mM 5HT administration (*level 1*: N = 7, *level 2*: N = 3). The number of the ants that responded with a turn response increased after 2hr (*level 0*: N = 3, *level 1*: N = 8, *level 2*: N = 8).

### Effects of oral administration of L-DOPA

To increase endogenous DA levels in the brain, its precursor L-DOPA was orally applied to the ants. Oral administration of 10mM L-DOPA solution significantly increased the defensiveness level score after 2hr (Fig. 4A). Before oral administration of 10mM L-DOPA, 60 workers were collected from 3 colonies (20 ants from each of 3 colonies) and their responses to the tactile stimulus observed. Fifty-three out of 60 ants responded with a dart response to the stimulus and they were given 10mM L-DOPA solution orally. Although no obvious behavioral change was observed after 1hr, the number of ants that showed a turn response increased to 19 out of 53 ants after 2hr (*level 1*: N = 11, *level 2*: N = 8). There were 9 ants that showed no obvious response to the stimulus (*level 0*).

**Fig. 4.**
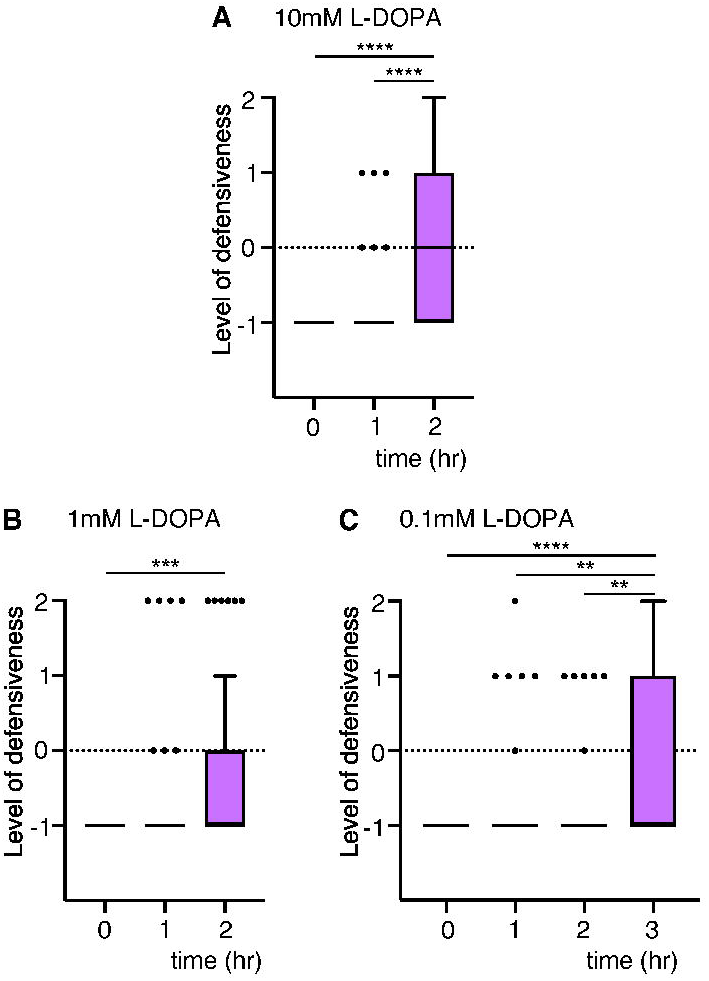
Pharmacological manipulation of endogenous DA in the brain using L-DOPA. Box-and-whisker graphs indicate minimum, median, maximum, 25% percentile and 75% percentile. **A:** Effect of oral administration of 10mM L-DOPA that is a precursor of DA. Defensiveness level scores did not change after 1hr, however significantly increased after 2hr. **B:** Effect of oral administration of 1mM L-DOPA. Defensiveness level scores did not change after 1hr but increased significantly after 2hr. **C:** Effect of oral application of 0.1mM L-DOPA. Defensiveness score levels did not change after 1hr and 2hr but increased significantly after 3hr. *: p < 0.05, **p< 0.01, ***: p < 0.001, ****: p< 0.0001

Oral administration of 1mM L-DOPA solution also significantly increased the defensiveness level score after 2hr (p=0.0007) (Fig. 4B). Sixty ants were randomly collected from 3 colonies (20 ants from each) and responses to the tactile stimulus prior to oral application of 1mM L-DOPA solution observed. Forty-seven out of 60 ants responded with a dart response to the tactile stimulus and were given 1mM L-DOPA solution orally. There was no obvious change in responses to the stimulus, but the number of ants that responded with a turn response increased after 2hr. Six out of 47 ants responded with a turn response (*level 1*: N = 3, *level 2*: N = 3) and 3 out of 47 ants ignored the stimulus (*level 0*). There was significant difference between 1mM L-DOPA and 10mM L-DOPA 2hr after oral administration (1mM L-DOPA 2hr vs. 10mM L-DOPA 2hr: p = 0.006).

Oral administration of 0.1mM L-DOPA significantly increased the defensiveness level score after 3hr (0hr vs. 3hr: p < 0.0001, 1hr vs. 3hr: p = 0.0036, 2hr vs 3hr: p = 0.0022) (Fig. 4C). Forty-five ants collected from 3 colonies (15 ants from each of 3 colonies) were used. Forty-one out of 45 ants showed a dart response to the stimulus before oral administration. There was no obvious effect on the responses to the tactile stimulus after 1hr and 2hr. However, the number of the ants that responded with a turn increased to 13 out of 41 ants after 3hr (*level 0*: N = 3, *level 1*: N = 9, *level 2*: N = 4). Note that there were not as many ants representing the level 2 defensive turn response to the tactile stimulus.

### Effects of oral administration of 5HTP

To increase endogenous 5HT in the ants, 5HTP was orally applied (Fig. 5). Oral administration of 10mM 5HTP significantly increased the defensiveness score level after 1hr (0hr vs. 1hr: p = 0.0014) (Fig. 5A). Forty workers were randomly collected (20 ants from each colony) and the response to the tactile stimulus before oral administration observed. Since all of them responded with a dart response, they were all given 10mM 5HTP. The number of ants that responded with a defensive turn to the stimulus increased to 11 out of 40 ants after 1hr (*level 1*: N = 6, *level 2*: N = 5). The defensiveness level score significantly increased after 90 min (0hr vs. 1.5hr: p < 0.0001, 1hr vs. 1.5hr: p = 0.0002). Twenty three out of 40 ants responded with a turn response to the stimulus after 90 min (*level 0*: 1, *level 1*: N = 4, *level 2*: N = 19).

**Fig. 5.**
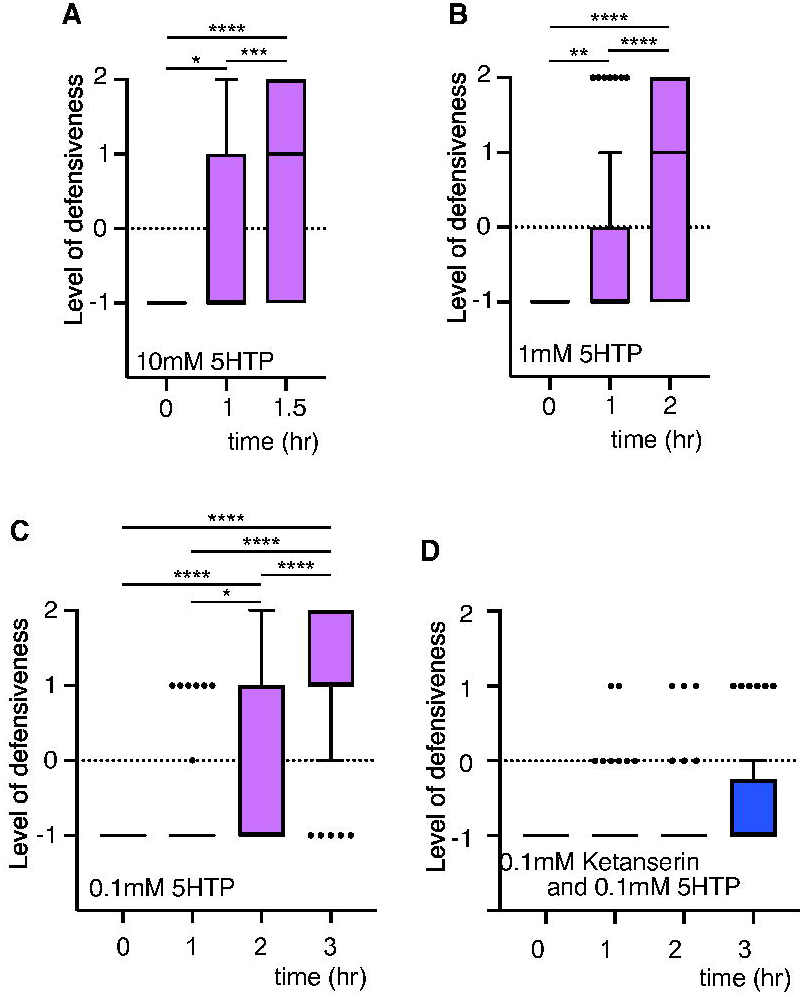
Effect of oral administration of 5HTP and ketanserin. Box-and-whisker graphs indicate minimum, median, maximum, 25% percentile and 75% percentile. **A:** Effect of oral administration of 10mM 5HTP (N = 20). Defensiveness level scores significantly increased after 1hr and 1.5hr. **B:** Effect of oral administration of 1mM 5HTP (N = 53). Defensiveness level scores increased after 1hr and 2hr. **C:** Effect of oral administration of 0.1mM 5HTP (N = 42). Defensiveness level scores did not change after 1hr. However, defensiveness level scores increased significantly after 2hr and 3hr. The effect of 0.1mM 5HTP was time dependent and the score at 3hr was significantly higher than that at 2hr. **D:** Effects of oral administration of a cocktail of 0.1mM ketanserin and 0.1mM 5HTP (N = 40). Oral administration of the cocktail did not change the defensiveness level score. Oral administration of the cocktail slightly increased the defensive level after 3hr, but it was not significant. *: p < 0.05, **p< 0.01, ***: p < 0.001, ****: p < 0.0001

The effect of 1 mM 5HTP on behavior was then examined. Oral administration of 1mM 5HTP significantly increased the defensiveness level score after 1hr (0hr vs. 1hr: p = 0.0086) and after 2hr (0hr vs. 2hr: p < 0.0001, 1hr vs. 2hr: p < 0.0001) (Fig.5B). Sixty ants were randomly collected from 3 colonies (20 ants from each colony) and responses to the tactile stimulus prior to oral application observed. Fifty-three out of 60 ants responded with a dart escape to the tactile stimulus and used for subsequent tests. Administration of 1mM 5HTP solution increased the number of ants that responded with a turn response to 9 out of 53 ants after 1hr (*level 1*: N = 4, *level 2*: N = 5). The number of ants that responded with a turn increased to 20 out of 53 after 2hr (*level 1*: N = 5, *level 2*: N = 15).

Oral administration of 0.1mM 5HTP did not increase the defensiveness score level after 1hr (Fig. 5C). However, it increased the defensiveness level score significantly after 2hr (p < 0.0001) and after 3hr (p < 0.0001). Sixty workers from 3 colonies were collected (20 ants each from each colony) and response to the tactile stimulus before administration of 5HTP observed. Forty-two ants responded with a dart response and used to examine the effect of 0.1mM 5HTP. The number of ants that responded with a turn response to the stimulus increased to 13 out of 42 ants after 2hr (*level 0*: N = 5, *level 1*: N = 11, *level 2*: N = 2). The number of the ants that responded with a turn response increased to 36 out of 42 ants after 3hr (*level 0*: N = 1, *level 1*: N = 24, *level 2*: N = 12). The effect of oral administration of 5HTP on initiating the turn response was both dose dependent and time dependent. Note that oral application of 5HTP increased the number of *level 2* defensive turns more than L-DOPA.

To confirm that increases in endogenous 5HT contribute to initiating defensive turn responses to the tactile stimulus, the effect of 5HT receptor antagonist ketanserin on behavior was examined (Fig. 5 D). Forty-five ants were randomly collected from 3 colonies (15 ants from each colony) and responses to the tactile stimulus prior to oral application observed. Forty out of 45 ants responded with a dart response to the tactile stimulus and given a cocktail of 0.1mM ketanserin and 0.1mM 5HTP diluted in a 20% sucrose solution orally. After 1 hr administration, most of the ants responded with a dart response (*level -1*: N = 32, *level 0*: N = 6, *level 1*: N = 2). After 2hr, most ants still responded with a dart response to the stimulus (*level -1*: N = 33, *level 0*: N = 3, *level 1*: N = 4). After 3hr, 6 out of 40 ants responded with a turn response (*level 1*) to the tactile stimulus (*level -1*: N = 30, *level 0*: N = 4, *level 1*: N = 6). There were no ants that responded with a turn with open mandible (*level 2*), when the cocktail of ketanserin and 5HTP was administered. This indicates that ketanserin inhibited the effect of 5HT that was generated by oral administration of 5HTP.

## DISCUSSION

An increase in aggressiveness during hunting and defensive behavior is closely linked to the generation of ultra-fast movements of the mandible in the trap-jaw ant. Many studies have focused on rapid movement of the trap-jaw i.e. its sensory-motor control and neuroanatomy (Gronenberg, 1995; Gronenberg, 1996b; Gronenberg et al., 1998; Gronenberg et al., 1993), kinetics (Patek et al., 2006) and ecological relevance (Larabee and Suarez, 2015). However, it remains unclear how the nervous system modulates aggressiveness to initiate defensive behavior in the trap-jaw ant. This study aimed to gain an understanding of the neuronal mechanism underlying the initiation of defensive movements.

The predominant response to unexpected touch in the trap-jaw ant was a dart escape. Less than 10% of workers in the colony made defensive turns. Aggressiveness is crucial not only to initiate defensive behavior against enemies but also to establish sociality in ants (Cuvillier-Hot et al., 2002) (Tanner and Adler, 2009). The ants that responded with defensive turns to touch showed antennal boxing. Antennal boxing could function to identify the source of stimulation and is associated with aggression during agonistic behavior and hunting in ponerine ants (Gobin et al., 1998). Few workers escalated their responses to violent attack against the drawing brush used for stimulation in this study, indicating that the ants might not identify the drawing brush as an urgent threat nor a prey. It is thought that the colony size of ponerine ants is associated with a hunting strategy (Beckers et al., 1989). Ponerine ants whose colony size are 200-300 individuals, such as *O. kuroiwae*, are thought to go hunting alone or in tandem. This suggests that social aggression in the ant *O. kuroiwae* is mostly suppressed and less than 10% of workers in a colony maintain aggressive potentially to play a role in foraging or grading of the colony.

Brain biogenic amines OA, DA and 5HT are candidate neuromodulators that regulate aggressiveness to initiate defensive behavior in the trap-jaw ant. The levels of these amines in the brain were significantly elevated in the ants that responded with defensive turns to the tactile stimulus compared to those in the ants that responded with dart escape to the stimulus (Fig. 2).

Oral administration of OA did not affect the initiation of defensive behavior in the trap-jaw ant. The OAergic system in the ant may be involved in behaviors other than initiating defensive behavior, although elevation of brain OA is associated with an increase in aggressiveness in insects (i.e. cricket: (Stevenson et al., 2005) (Rillich et al., 2011b), *Drosophila*: (Zhou et al., 2008), and the ant *Formica japonica*, (Aonuma and Watanabe, 2012b). The actions of the OAergic system and TAergic system in insects are thought to be homologous with the noradrenergic system in vertebrates (Roeder, 1999). OA acts as a multifunctional mediator in insects. OA, and its precursor TA, itself mediates defensive behavior of soldiers in termites (Ishikawa et al., 2016). Brain OA increases pheromone sensitivity in the silkmoth (Pophof, 2000, 2002; Gatellier et al., 2004). The OAergic system in the brains of social insects is associated with nest-mate recognition (honeybee: (Robinson et al., 1999); fire ant: (Vander Meer et al., 2008)). It has also been demonstrated that OA functions to generate energy for muscle activity via activation of trehalase (Candy, 1978; Jahagirdar et al., 1984; Vaandrager et al., 1988; Orchard et al., 1993). During predator-prey encounters, the trap-jaw ant expresses defensive or threatening postures that include opening the mandibles and antennal boxing. The mandibles have sensory hairs to detect prey or predators and close extremely rapidly (Gronenberg, 1995; Gronenberg et al., 1993). Furthermore, strong contractions of the adductor muscles before closing the mandible contribute to generate the ultra-fast movement. Elevation of OA in the brain could be associated with increasing sensitivity of chemotactile signals and/or muscle metabolism to generate the ultra-fast movement of the mandibles. Further studies to investigate the roles of OA in the trap-jaw ant are necessary to determine the precise role that OA plays in aggressive behavior.

Oral administration of DA and its precursor L-DOPA increased the initiation of turn responses to unexpected touch, although they rarely opened their mandible. This suggests that the DAergic system is weakly involved in the initiation of defensive responses to unexpected tactile stimuli. It has been demonstrated that a pharmacological increase in DA levels elevates aggressiveness towards prey in the ant *Formica polyctena* (Szczuka et al., 2013). The DAergic system is closely associated with nest-mate hierarchy and ovarian activity in the ponerine ant *H. saltator* (Penick et al., 2014). The results of this study support the modulatory role of DA on elevation of aggressiveness.

The DAergic system is thought to be multifunctional. DA is closely associated with reproductive behavior in insects (Gruntenko et al., 2005; Sasaki and Harano, 2010). It has been shown that increases in the levels of DA in the brain are associated with the initiation of egg-laying behavior in the eusocial wasp (Sasaki et al., 2007). DA is thought to linked with copulation behavior in female *Drosophila*, (Neckameyer, 1998). In the ant *Diacamma* sp. the workers decrease the amount DA in the brain by making contact with the queen, which in turn suppresses their aggressiveness towards their nest mates (Shimoji et al., 2017). In honeybees, the mandibular gland pheromone of the queen modulates the action of DA in the brain of workers (Beggs et al., 2007; Beggs and Mercer, 2009) and increases in the levels of DA in the brains of workers is closely associated with development of their ovary (Sasaki and Nagao, 2001). These previous findings suggest that there is only a weak involvement of the DAergic system in defensive behavior of the trap-jaw ant but instead serves to avoid activating the reproduction of workers and to avoid conflicts among nest-mates.

Oral administration of 5HT and its precursor 5HTP increased the number of defensive turn responses to unexpected tactile stimulation in the trap-jaw ant. It is believed that 5HT contributes to the modulation of aggressive behavior in invertebrates (Kravitz and Huber, 2003) as well as vertebrates (Olivier, 2004; Montoya et al., 2012). This study has demonstrated that by contrast to the administration of DA and L-DOPA, the ants opened the mandibles widely and showed antennal boxing. Furthermore, administration of ketanserin, an inhibitor of the 5HT_2_ receptor, inhibited the effects of 5HTP, which indicates that endogenous 5HT release modulates the initiation of defensive behavior in trap-jaw ants. It was previously shown that a tonic increase in 5HT_2_ receptor escalates aggression, and ketanserin strikingly abolishes aggressive behavior in mice (Shih et al., 1999; Takahashi et al., 2011). The 5HTergic system in the trap-jaw ant could be similar to the mammalian system in regulating aggressive behavior. 5HT in the brain has been shown to regulate aggressiveness in invertebrates. It was reported that elevation of brain 5HT is closely linked to aggression between interspecies and intraspecies in the ant *Formica rufa* (Tarchalska et al., 1975). Elevation of 5HT in the brain enhances aggressive behavior in *Drosophila* (Dierick and Greenspan, 2007). Oral administration of 5HTP and 5HT enhances the expression of high-intensity aggressive behaviors and increases the wining probability of agonistic contests in the stalk-eyed fly *Teleopsis dalmanni* (Bubak et al., 2014), and in crayfish 5HT increases aggressiveness of subordinates (Huber et al., 1997; Kravitz, 2000). The results of this study support these previous studies. On the other hand, the opposite effects of 5HT on aggressiveness in insects has also been reported. The serotonin 5-HT_2_ receptor suppresses aggressive behavior in *Drosophila* (Johnson et al., 2009). It has been demonstrated that 5HT depresses aggressiveness in subordinate crickets after agonistic interactions. (Rillich and Stevenson, 2018). It has also demonstrated that synaptic responsiveness to 5HT changes with social status in crayfish (Yeh et al., 1997). Social interactions are one of the important factors that maintains homeostasis of aminergic control in ants (Wada-Katsumata et al., 2011). Contact with nest-mate workers rescued depressed DA and OA levels in the ant. It is demonstrated that the sting alarm pheromone, isoamyl acetate upregulates brain 5HT and DA, which elevate workers’ aggressiveness to enhance social defensive behavior in honeybees (Bubak et al., 2020). Social insects change the titer of brain amines according to social experience. It is clearly important that we unveil the co-effects of brain amines to understand aggressiveness and the initiation of defensive behavior in the trap-jaw ant.

This study concludes that increases in brain 5HT levels elevate aggressiveness to initiate defensive responses to unexpected tactile stimuli and that brain DA weakly contributes to the initiate of defensive behavior in the trap-jaw ant. Further investigation of the co-effects of 5HT and DA on initiating defensive behavior in the trap-jaw ant would help us to unveil the neuronal mechanisms underlying social escape and social defense in social insects.

## Acknowledgements

I thank Prof. Philip L. Newland for his critical comments upon my manuscript. I also thank Dr. Hiroyuki Shimoji for his help to collect ants and thank Dr. Daiki Wakita, Mr. Keisuke Naniwa and Mrs. Yuka Hoshino for their comments and technical help. A part of this work was supported by grants-in-aid for JST CREST (Grant Number JPMJCR14D5) and JSPS KAKENHI (Grant-in-Aid for Scientific Research (S), Grant Number JP17H06150), Japan.

## Author contribution

H.A. conceived and designed the experiment; H.A. performed the experiment and analyzed the data; H.A. wrote the paper.

## Supplementary Information

**S1.** Hunting behavior of the trap-jaw ant *Odontomachus kuroiwae*. The ant orients to a small insect while opening the mandibles widely. After it detects and identifies the prey using the antennae, the mandibles close extremely fast, to bite, and then it stings to inject venom into the prey.

**S2.** Behavioral responses of the trap-jaw ant to unexpected tactile stimuli to the abdomen. The movie indicates the *Level -1* dart response, *Level 0* no response, *Level 1* turn response without open mandibles, and *Level 2* turn response with open mandibles.

**S3.** Amount of precursor and catabolites of biogenic amines in the brain. Box-and-whisker graphs indicate minimum, median, maximum, 25% percentile and 75% percentile. A: Amount of 5HTP in the brain. There was no significant difference between turn and dart (Unpaired t test with Welch’s correction: p = 0.21). B: Amount of Nac-5HT in the brain. There was no significantly difference between turn and dart (Unpaired t test with Welch’s correction: p = 0.18). C: Amount of Nac-DA in the brain. There was no significant difference between turn and dart (Unpaired t test with Welch’s correction: p = 0.80). D: Amount of Nac-TA in the brain. There was no significant difference between turn and dart (Unpaired t test with Welch’s correction: p = 0.75).

## REFERENCES

Alexander, D. R. (1961). Aggressiveness, territoriality, and sexual behavior in field crickets (Orthoptera: Gryllidae). Behaviour 17, 130–223.

Aonuma, H., Nagayama, T. and Hisada, M. (1994). Output effect of identified ascending interneurons upon the abdominal postural system in the crayfish *Procambarus clarkii* (Girard). Zoolog Sci 11, 191–202.

Aonuma, H. and Watanabe, T. (2012a). Changes in the content of brain biogenic amine associated with early colony establishment in the Queen of the ant, Formica japonica. PLoS ONE 7, e43377.

Aonuma, H. and Watanabe, T. (2012b). Octopaminergic system in the brain controls aggressive motivation in the ant, Formica japonica. Acta Biol Hung 63 Suppl 2, 63–8.

Baumann, A., Blenau, W. and Erber, J. (2003). Biogenic amines In i (eds. Resh VH and Cradè RT). San Diego, California. USA: Academic Press., p. 91–94.

Beckers, R., Goss, S., Deneubourg, J.-L. and Pasteels, J.-M. (1989). Colony size, communication and ant foraging strategy. Psyche: A Journal of Entomology 96, 239–256.

Beggs, K. T., Glendining, K. A., Marechal, N. M., Vergoz, V., Nakamura, I., Slessor, K. N. and Mercer, A. R. (2007). Queen pheromone modulates brain dopamine function in worker honey bees. Proceedings of the National Academy of Sciences 104, 2460–2464.

Beggs, K. T. and Mercer, A. R. (2009). Dopamine receptor activation by honey bee queen pheromone. Curr Biol 19, 1206–9.

Bubak, A. N., Renner, K. J. and Swallow, J. G. (2014). Heightened serotonin influences contest outcome and enhances expression of high-intensity aggressive behaviors. Behav Brain Res 259, 137–142.

Bubak, A. N., Watt, M. J., Yaeger, J. D., Renner, K. J. and Swallow, J. G. (2020). The stalk-eyed fly as a model for aggression–is there a conserved role for 5-HT between vertebrates and invertebrates? Journal of Experimental Biology 223.

Candy, D. J. (1978). The regulation of locust flight muscle metabolism by octopamine and other compounds. Insect Biochemistry 8, 177–181.

Choe, J. C. and Crespi, B. J. (1997). Morphologically “primitive” ants: Comparative review of social characters, and the importance of queen-worker dimorphism. The evolution of social behaviour in insects and arachnids: Cambridge University Press.

Cuvillier-Hot, V., Gadagkar, R., Peeters, C. and Cobb, M. (2002). Regulation of reproduction in a queenless ant: aggression, pheromones and reduction in conflict. Proceedings of the Royal Society of London. Series B: Biological Sciences 269, 1295–1300.

De la Mora, A., Pérez-Lachaud, G. and Lachaud, J.-P. (2008). Mandible strike: the lethal weapon of Odontomachus opaciventris against small prey. Behavioural processes 78, 64–75.

Dierick, H. A. and Greenspan, R. J. (2007). Serotonin and neuropeptide F have opposite modulatory effects on fly aggression. Nat Genet 39, 678–682.

Edwards, D. H. and Kravitzt, E. A. (1997). Serotonin, social status and aggression. Current Opinion in Neurobiology 7, 812–817.

Evans, P. C. (1980). Biogenic amines in the insect nervous system. Adv. Insect physiol. 422, 317–322.

Gobin, B., Billen, J. and Peeters, C. (2001). Dominance interactions regulate worker mating in the polygynous ponerine ant Gnamptogenys menadensis. Ethology 107, 495–508.

Gobin, B., Peeters, C., Billen, J. and Morgan, E. D. (1998). Interspecific trail following and commensalism between the ponerine ant Gnamptogenys menadensis and the formicine ant Polyrhachis rufipes. Journal of Insect Behavior 11, 361–369.

Gronenberg, W. (1995). The fast mandible strike in the trap-jaw ant Odontomachus. J Comp Physiol A 176, 399–408.

Gronenberg, W. (1996a). Neuroethology of ants. Naturwissenschaften 83, 15–27.

Gronenberg, W. (1996b). The trap-jaw mechanism in the dacetine ants Daceton armigerum and Strumigenys sp. J Exp Biol 199, 2021–33.

Gronenberg, W., Hölldobler, B. and Alpert, G. D. (1998). Jaws that snap: control of mandible movements in the ant Mystrium. Journal of insect physiology 44, 241–253.

Gronenberg, W., Tautz, J. and Hölldobler, B. (1993). Fast trap jaws and giant neurons in the ant Odontomachus. Science 262, 561–563.

Gruntenko, N., Karpova, E., Adonyeva, N., Chentsova, N., Faddeeva, N., Alekseev, A. and Rauschenbach, I. Y. (2005). Juvenile hormone, 20-hydroxyecdysone and dopamine interaction in Drosophila virilis reproduction under normal and nutritional stress conditions. Journal of insect physiology 51, 417–425.

Holway, D. A., Suarez, A. V. and Case, T. J. (1998). Loss of intraspecific aggression in the success of a widespread invasive social insect. Science 282, 949–952.

Hoyer, S. C., Eckart, A., Herrel, A., Zars, T., Fischer, S. A., Hardie, S. L. and Heisenberg, M. (2008). Octopamine in male aggression of Drosophila. Curr Biol 18, 159–67.

Huber, R., Smith, K., Delago, A., Isaksson, K. and Kravitz, E. A. (1997). Serotonin and aggressive motivation in crustaceans: altering the decision to retreat. Proc Natl Acad Sci U S A 94, 5939–42.

Ishikawa, Y., Aonuma, H., Sasaki, K. and Miura, T. (2016). Tyraminergic and Octopaminergic Modulation of Defensive Behavior in Termite Soldier. PLoS ONE 11, e0154230.

Jahagirdar, A., Downer, R. and Viswanatha, T. (1984). Influence of octopamine or trehalase activity in muscle and hemolymph of the American cockroach, Periplaneta americana L. Biochimica et Biophysica Acta (BBA)-General Subjects 801, 177–183.

Johnson, O., Becnel, J. and Nichols, C. D. (2009). Serotonin 5-HT(2) and 5-HT(1A)-like receptors differentially modulate aggressive behaviors in Drosophila melanogaster. Neuroscience 158, 1292–300.

Just, S. and Gronenberg, W. (1999). The control of mandible movements in the ant Odontomachus. J Insect Physiol 45, 231–240.

Kostowski, W., Tarchalska, B. and Wanchowicz, B. (1975). Brain catecholamines, spontaneous bioelectrical activity and aggressive behavior in ants (Formica rufa). Pharmacol Biochem Behav 3, 337–42.

Kravitz, E. (2000). Serotonin and aggression: insights gained from a lobster model system and speculations on the role of amine neurons in a complex behavior. J Comp Physiol A 186, 221–238.

Kravitz, E. A. and Huber, R. (2003). Aggression in invertebrates. Current Opinion in Neurobiology 13, 736–743.

Larabee, F. J. and Suarez, A. V. (2015). Mandible-Powered Escape Jumps in Trap-Jaw Ants Increase Survival Rates during Predator-Prey Encounters. PLoS ONE 10, e0124871.

Montoya, E. R., Terburg, D., Bos, P. A. and Van Honk, J. (2012). Testosterone, cortisol, and serotonin as key regulators of social aggression: A review and theoretical perspective. Motivation and emotion 36, 65–73.

Nagayama, T., Takahata, M. and Hisada, M. (1986). Behavioral transition of crayfish avoidance reaction in response to uropod stimulation. Experimental biology 46, 75–82.

Neckameyer, W. S. (1998). Dopamine and mushroom bodies in Drosophila: experience-dependent and -independent aspects of sexual behavior. Learn Mem 5, 157–65.

Ohkawara, K. and Aonuma, H. (2016). Changes in the levels of biogenic amines associated with aggressive behavior of queen in the social parasite ant Vollenhovia nipponica. Insectes Sociaux 63, 257–264.

Olivier, B. (2004). Serotonin and aggression. Annals of the New York Academy of Sciences 1036, 382–392.

Orchard, I., Ramirez, J.-M. and Lange, A. B. (1993). A multifunctional role for octopamine in locust flight. Annual review of entomology 38, 227–249.

Patek, S., Baio, J., Fisher, B. and Suarez, A. (2006). Multifunctionality and mechanical origins: ballistic jaw propulsion in trap-jaw ants. Proceedings of the National Academy of Sciences 103, 12787–12792.

Penick, C. A., Brent, C. S., Dolezal, K. and Liebig, J. (2014). Neurohormonal changes associated with ritualized combat and the formation of a reproductive hierarchy in the ant Harpegnathos saltator. J Exp Biol 217, 1496–503.

Rillich, J., Schildberger, K. and Stevenson, P. A. (2011a). Octopamine and occupancy: an aminergic mechanism for intruder-resident aggression in crickets. Proc. R. Soc. B 278, 1873–1880.

Rillich, J., Schildberger, K. and Stevenson, P. A. (2011b). Octopamine and occupancy: an aminergic mechanism for intruder–resident aggression in crickets. Proceedings of the Royal Society B: Biological Sciences 278, 1873–1880.

Rillich, J. and Stevenson, P. A. (2018). Serotonin mediates depression of aggression after acute and chronic social defeat stress in a model insect. Front Behav Neurosci 12, 233.

Robinson, G. E., Heuser, L. M., LeConte, Y., Lenquette, F. and Hollingworth, R. M. (1999). Neurochemicals aid bee nestmate recognition. Nature 399, 534–535.

Roeder, T. (1999). Octopamine in invertebrates. Prog Neurobiol 59, 533–61.

Roeder, T. (2005). Tyramine and octopamine: ruling behavior and metabolism. Annu Rev Entomol 50, 447–77.

Sasaki, K. and Harano, K.-i. (2010). Multiple regulatory roles of dopamine in behavior and reproduction of social insects. Trends Entomol 6, 1–13.

Sasaki, K. and Nagao, T. (2001). Distribution and levels of dopamine and its metabolites in brains of reproductive workers in honeybees. Journal of insect physiology 47, 1205–1216.

Sasaki, K., Yamasaki, K. and Nagao, T. (2007). Neuro-endocrine correlates of ovarian development and egg-laying behaviors in the primitively eusocial wasp (Polistes chinensis). Journal of insect physiology 53, 940–949.

Seid, M. A. and Traniello, J. F. A. (2005). Age-related changes in biogenic amines in individual brains of the ant *Pheidole dentata*. Naturwissenschaften 92, 198–201.

Seid, M. A. and Traniello, J. F. A. (2006). Age-related repertoire expansion and division of labor in Pheidole dentata *(Hymenoptera: Formicidae):* a new perspective on temporal polyethism and behavioral plasticity in ants. Behav Ecol Sociobiol 60, 631–644.

Shih, J. C., Ridd, M. J., Chen, K., Meehan, W. P., Kung, M.-P., Seif, I. and De Maeyer, E. (1999). Ketanserin and tetrabenazine abolish aggression in mice lacking monoamine oxidase A. Brain Res 835, 104–112.

Shimoji, H., Aonuma, H., Miura, T., Tsuji, K., Sasaki, K. and Okada, Y. (2017). Queen contact and among-worker interactions dually suppress worker brain dopamine as a potential regulator of reproduction in an ant. Behavioral Ecology and Sociobiology 71.

Song, C.-K., Herberholz, J. and Edwards, D. H. (2006). The effects of social experience on the behavioral response to unexpected touch in crayfish. Journal of Experimental Biology 209, 1355–1363.

Stevenson, P. A., Dyakonova, V., Rillich, J. and Schildberger, K. (2005). Octopamine and experience-dependent modulation of aggression in crickets. J Neurosci 25, 1431–41.

Szczuka, A., Korczyńska, J., Wnuk, A., Symonowicz, B., Szwacka, A. G., Mazurkiewicz, P., Kostowski, W. and Godzińska, E. J. (2013). The effects of serotonin, dopamine, octopamine and tyramine on behavior of workers of the ant Formica polyctena during dyadic aggression tests. Acta Neurobiol Exp 73, 495–520.

Takahashi, A., Quadros, I. M., de Almeida, R. M. and Miczek, K. A. (2011). Brain serotonin receptors and transporters: initiation vs. termination of escalated aggression. Psychopharmacology (Berl) 213, 183–212.

Tanner, C. J. and Adler, F. R. (2009). To fight or not to fight: context-dependent interspecific aggression in competing ants. Animal Behaviour 77, 297–305.

Tarchalska, B., Kostowski, W., Markowska, L. and Markiewicz, L. (1975). On the role of serotonin in aggressive behaviour of ants Genus formica. Pol J Pharmacol Pharm 27, 237–9.

Tomioka, K., Ikeda, M., Nagao, T. and Tamotsu, S. (1993). Involvement of serotonin in the circadian rhythm of an insect visual system. Naturwissenschaften 80, 137–139.

Vaandrager, S., Wynne, H. and Beenakkers, A. T. (1988). Regulation of flight related trehalose utilization in the locust Locusta migratoria. Comparative Biochemistry and Physiology Part A: Physiology 91, 653–657.

Vander Meer, R. K., Preston, C. A. and Hefetz, A. (2008). Queen regulates biogenic amine level and nestmate recognition in workers of the fire ant, Solenopsis invicta. Naturwissenschaften 95, 1155–8.

Vleugels, R., Verlinden, H. and Broeck, J. V. (2015). Serotonin, serotonin receptors and their actions in insects. Neurotransmitter 2, e314.

Wada-Katsumata, A., Yamaoka, R. and Aonuma, H. (2011). Social interactions influence dopamine and octopamine homeostasis in the brain of the ant *Formica japonica*. J Exp Biol 214, 1707–13.

Yakovlev, I. (2018). Effects of octopamine on aggressive behavior in red wood ants. Neuroscience and Behavioral Physiology 48, 279–288.

Yeh, S. R., Musolf, B. E. and Edwards, D. H. (1997). Neuronal adaptations to changes in the social dominance status of crayfish. J Neurosci 17, 697–708.

Zhou, C., Rao, Y. and Rao, Y. (2008). A subset of octopaminergic neurons are important for Drosophila aggression. Nat Neurosci 11, 1059.

