## Supplementary figures and images for "Serotonergic control in initiating defensive responses to unexpected tactile stimuli in the trap-jaw ant *Odontomachus kuroiwae*"

### S3

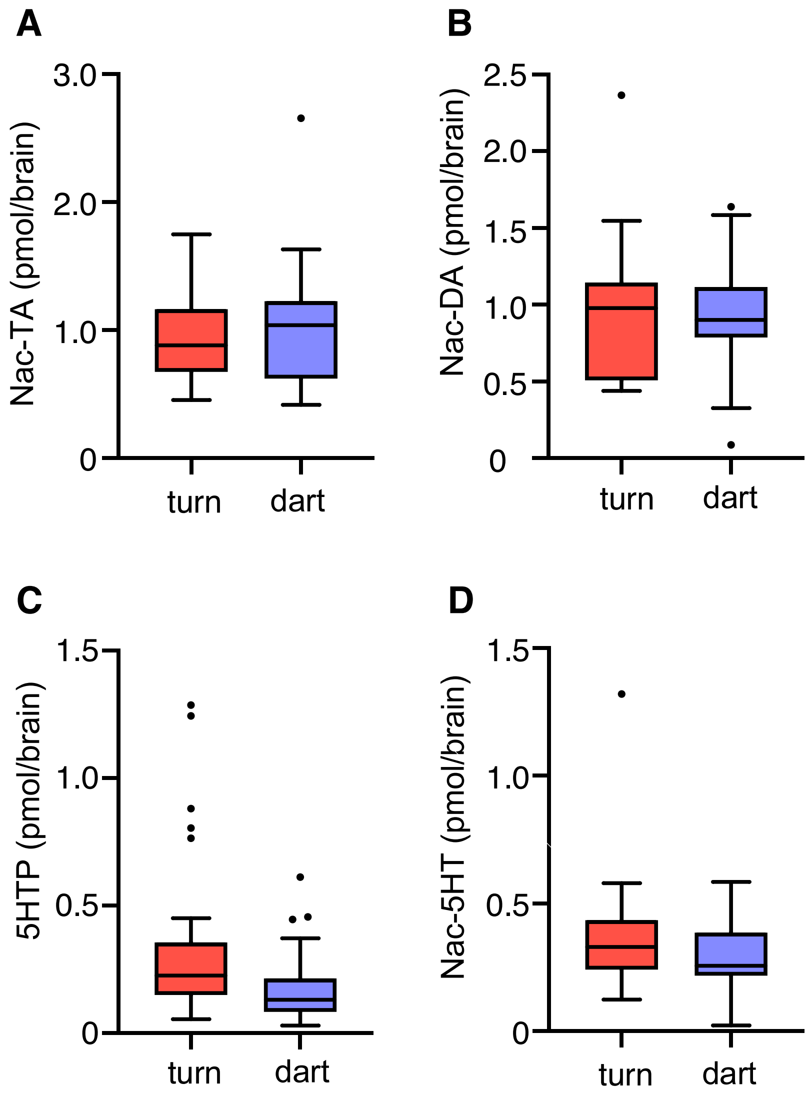
